# Misleading monitoring: more hatchlings do not represent turtle population recovery

**DOI:** 10.1101/2025.02.14.638251

**Authors:** Darren Norris

## Abstract

This article critiques a recent study by Lacava et al. (2024) that suggests the population of *Podocnemis expansa* has recovered in Brazil based on increased hatchling releases across 11 nesting areas. I present three main challenges to the study conclusions. Firstly, hatchling counts alone should not be used to assess population recovery. Secondly, the study failed to consider alternative hypotheses that could explain the observed increases, including changes in methods and effort over time. Lastly, the statistical analysis was flawed, specifically neglecting temporal autocorrelation and failing to distinguish between statistical significance and biological relevance. Reanalysis of the original data, accounting for temporal autocorrelation, revealed sustained increases in hatchlings released in only five of eleven areas, contradicting the original claim of increases in most areas. Furthermore, reanalysis suggests that methodological changes influenced results in at least one area. The study published by Lacava et al. (2024) requires correction to avoid ingenuous and potentially misleading conclusions and to more closely reflect the reality of 21^st^-century freshwater turtle conservation efforts.

---

Recently, Lacava et al. (2024) showed that the number of *Podocnemis expansa* hatchlings released had increased across nesting beaches over four decades. The authors showed that currently the Brazilian Amazon Chelonian Program (Programa Quelônios da Amazônia - PQA) typically releases more than two million hatchlings per year. This number is truly impressive, and the scale of the actions represent a significant conservation success. While the study presents a positive trend in hatchlings released in some nesting areas, the data collected and the results presented do not support the authors’ conclusions regarding population recovery. Reanalysis including temporal autocorrelation showed evidence of a sustained increase in hatchlings released at only five of the 11 areas (Supplemental Material), which challenges the authors’ conclusion of increases in “most” areas. Here I identify where corrections are required, grouped into three general themes: i) hatchling counts alone do not represent turtle population trends, ii) authors fail to consider equally plausible alternative hypotheses, and iii) incorrect use of statistical significance.

Firstly, the authors use annual hatchling counts in isolation to evaluate population trends. While authors identify hatchling counts as the most consistently collected data available, this does not mean it can be used to assess population trends. Studies with *P. expansa* and sea turtle species with similar nesting patterns demonstrated that while hatchling counts can be part of turtle population monitoring, they’re inadequate in isolation (Eisemberg et al. 2019; Forero-Medina et al. 2021; Hays et al. 2025; Hendrix and Pérez-Espona 2024). A more appropriate approach would be to simply refer to hatchlings released instead of population trends/recovery. The potential of hatchlings released to reflect population trends could then perhaps be discussed. For example, reinforcing limitations and the need to include other factors like adult survival for assessing turtle population trends (Gibbs and Amato 2000; Páez et al. 2015).

Turtle population dynamics are strongly dependent on adult survival, as such indicators are required that are in some way calibrated with adult populations (Eisemberg et al. 2019; Moll and Moll 2004; Páez et al. 2015). The failure to include such indicators (e.g. hatchlings per nest/hatchlings per female) invalidates the authors’ conclusions regarding population trends. As reinforced by Eisemberg et al. (2019), the number of hatchlings released is a poor indicator of *P. expansa* population trends due to a dependence on environmental and climate factors (e.g. temperature and river levels). In the case of protected areas such as in Lacava et al. (2024), the number of hatchlings released will also depend on the conservation management actions and efforts (techniques, number of staff, etc.).

Before drawing any conclusions about trends based on hatchling counts, key assumptions need to be more rigorously tested. By emphasizing population recovery, the study assumes that an increased number of hatchlings released translates to increases in the adult breeding population. This assumption needs to be tested, for example by conducting post-release monitoring to assess actual hatchling survival rates in the wild (Mogollones et al. 2010). To address the limitations of using hatchling counts in isolation, the study could be refocused to assess changes in hatchling relative abundance over time, without drawing conclusions regarding population trends. By refocusing the study in this way, the authors could still provide valuable information about the impact of the PQA on hatchling production, while avoiding overinterpretation of the results.

Secondly, several equally plausible alternative hypotheses could also explain the observed trends. These hypotheses should be tested to avoid pitfalls in simplistic monitoring (Christie et al. 2021; Christie et al. 2019; Wauchope et al. 2021), and include: i) improved methods/techniques in nest and hatchling management over time leading to increases compared with the early years/decades, ii) increased number of staff/workers, iii) improved technical experience/capacity of staff/workers, iv) improved data collection and reporting, and v) expansion in the size/number of nesting areas included in conservation actions over the years, etc. As authors failed to provide key methodological details it is impossible to assess the validity of the methods. There is also no indication whether management efforts changed over time. For example, there is no detail on when the fencing of the hatchlings started in the different areas, the annual coverage of the fenced areas at the nesting beaches, or if the coverage changed over time. Indeed, it is unclear what was actually measured as in the methods authors state “the number of hatchlings released annually at each reproductive site”, whereas the legend for Figure 2 states: “Fig. 2 The number of Giant South American River Turtle (*Podocnemis expansa*) hatchlings born”. Without testing these alternative hypotheses, it is impossible to separate real trends from artifacts and systematic biases caused by variations in monitoring/management intensity and techniques.

Authors suggested in the discussion that the more positive trends they found in comparison to earlier studies were explained by the inclusion of more years: “Based on a more extended data set than previous studies on the Brazilian PQA public dataset (Eisemberg et al. 2019; Forero-Medina et al. 2019), our results show a far more positive trend of increase over 43 years in most areas and for all the nesting sites pooled together.”. This is not the case. Running the same analysis using subsets corresponding to the years in earlier studies, the trends can worsen or improve over time at different beaches (Supplemental Material S1). The contrasting results presented by the authors most likely came from the far more robust analysis presented in the earlier studies. The earlier studies used a combination of indicators not only hatchling counts and attempted to correct methodological differences over time (Eisemberg et al. 2019; Forero-Medina et al. 2021). The more cautious conclusions presented in the earlier studies therefore appear to have much stronger support based on the available data. It is unclear why Lacava et al. (2024) did not repeat the same analysis as presented in the earlier studies. One possible explanation is that a lack of robust monitoring data comes from severe budgetary cuts and limited operational cover, which limits the scope and activities of the PQA (Eisemberg et al. 2019).

Finally, there is an equivocal interpretation of statistical significance, with the authors also failing to appropriately address the biological relevance of the results. The authors used Generalized Linear Models (GLMs) to identify increases in hatchlings released at six of the 11 areas. This proportion means that overall, there was an equal chance for an area to experience increases or decreases in the number of hatchlings released (Chi-squared = 0, df = 1, p-value = 1). Statements such as “efforts for turtle conservation over four decades to protect the *P. expansa* were efficient in most PQA-protected nesting areas”, are therefore potentially misleading. As the study had no control areas, i.e. areas without protection, it is impossible to know if this overall proportion provides evidence of conservation success. In other words, the simple study design cannot be used to assess conservation effectiveness (Christie et al. 2021; Christie et al. 2019; Wauchope et al. 2021).

The six areas where “populations increased over time with statistical significance” were Abufari and Embaubal (“more subtle increase”), and Monte Cristo, Guaporé, Camaleões, and Walter Bury with a “very evident increase of hatchlings released over the 40 years.”. Both areas with “more subtle increase” had very low model deviance explained (12.9 and 18.2 % model deviance explained for Abufari and Embaubal respectively), with the lower confidence limits at the end of the monitoring period appearing to overlap the upper confidence limits at the start in both cases (Figure 2 in the article). This overlap in confidence intervals raises questions regarding statistical significance (Greenland et al. 2016; Halsey 2019; Muff et al. 2022).

As the analysis was based on counts over time, temporal autocorrelation needed to be explicitly included in the models to avoid type I errors (Carroll and Pearson 2000; Keith et al. 2015; Zuur et al. 2010). Reanalysis of the data including temporal autocorrelation with Generalized Linear Mixed Models provided evidence of increased hatchlings from five of the 11 nesting areas (Supplemental Material S1). Abufari beach was no longer “statistically significant”. The “significant increase” at this beach was due to a jump in the number of hatchlings released around 1990. Until 1990 there was an average of approximately 50 thousand hatchlings released, then after 1990 the number suddenly quadrupled to approximately 200,000 hatchlings (Figure 2 in the article). Such a sudden and large jump is not consistent with the population dynamics of turtle species. There has also been a steady decline in the number of hatchlings released at Abufari since 2005. These changes would not be reflected by the simplistic GLM approach adopted by the authors. The most likely explanation for the jump is a change in management such as increasing the area protected, change in counting technique, etc. This example highlights that a lack of methodological details makes it impossible to know how much of the observed increases are due to methodological differences or biologically meaningful changes in the numbers of hatchlings released.

How the numbers of hatchlings released translates into meaningful conservation results depends among other things on the biological relevance of the statistical significance. For example, while Walter Bury was the area with the strongest increase in hatchlings released, interviews with local communities around this area identified the sale of adults as the main reason for declines in turtle populations in the region (de Paula Nascimento et al. 2024). The same study identified a lack of ethics among beach guards as a problem around this area. Indeed, locals report beach guards selling adult turtles to middle-class buyers (de Paula Nascimento et al. 2024). This contrast between the increasing number of hatchlings and reports of reduced adult populations supports previous studies that highlighted a lack of correlation between hatchling counts and adult populations and reinforces an urgent need to redirect conservation efforts (Eisemberg et al. 2019; Páez et al. 2015). Recent declines over the past decade in another area (Guapore) were not represented by the GLM model chosen by the authors (Figure 2 in the published article). These declines are thought to be associated at least in part with extreme climatic events (Energisa 2024), which highlights how sensitive hatchling counts are to external factors and supports the need to not only appropriately quantify environmental and methodological differences among beaches and over time, but to also expand monitoring to include different stages including nests and adult females (Eisemberg et al. 2019; Forero-Medina et al. 2021; Páez et al. 2015).

I grouped corrections into three different sections, yet there are important considerations that can be found across all. The activities of PQA remain strongly focused on protecting beaches without considering the adult population. Despite calls for changes, the monitoring adopted by the PQA does not appear to have evolved to meet 21^st^-century challenges of rapid expansion of human-induced impacts across Amazonia, including increased extreme climatic events and habitat loss (Eisemberg et al. 2019; Páez et al. 2015). While quantifying absolute values remains challenging, there are a variety of cost-effective techniques that could be adopted to enable robust monitoring of population trends. An integrated approach is required that leverages the power of citizen science and digital technology to create a cost-effective and sustainable population monitoring program for *P. expansa*. Population monitoring could be enabled by integrating and calibrating techniques such as nest monitoring, mark-recapture and interviews across a network of community-based monitoring sites (Hallwass et al. 2013; Jones et al. 2008; Lahoz-Monfort and Magrath 2021). Such approaches have already been established and applied for monitoring sea turtles (Hays et al. 2025; Hendrix and Pérez-Espona 2024; Piacenza et al. 2019).

Without adapting the monitoring protocols to include nests and adult females it remains impossible to assess *P. expansa* population trends or conservation actions. Authors showed that there are typically more than two million hatchlings released per year. Unfortunately, considering what is known of the species population demographics, this is still well below the number required to replace individuals consumed annually in urban and rural areas in the Brazilian Amazon (Chaves et al. 2021; Peres 2000). Using what is known about turtle population dynamics, it is likely that 200 – 1000 hatchlings are needed to replace 1 adult for this long-lived and late-reproducing species. Although there is no published estimate covering the whole Brazilian Amazon, based on published data (Chaves et al. 2021; Peres 2000) it is plausible that somewhere around 300,000 adult *P. expansa* are consumed annually in Brazil – therefore at least 60 million hatchlings need to be released per year to replace the adults consumed.

In conclusion, while hatchling counts can be part of turtle population monitoring, they’re inadequate on their own to assess the effectiveness of conservation actions. Turtle species require not only long-term but also multi-faceted integrated approaches to both assess population trends and to guide and evaluate conservation efforts. The study published by Lacava et al. (2024) requires correction/clarification to avoid ingenuous and potentially misleading conclusions and to more closely reflect the reality of 21^st^-century freshwater turtle conservation efforts.

## Notes

### Competing Interest Statement

The authors have declared no competing interest.

### Summary of Updates

Improve text flow. Added examples from sea turtle monitoring. Corrected minor typos in the text.

## References

Carroll, SS, Pearson, DL 2000. Detecting and Modeling Spatial and Temporal Dependence in Conservation Biology. Conserv. Biol. 14: 1893–1897. 10.1111/j.1523-1739.2000.99432.x.

Chaves, WA, Valle, D, Tavares, AS, Morcatty, TQ, Wilcove, DS 2021. Impacts of rural to urban migration, urbanization, and generational change on consumption of wild animals in the Amazon. Conserv. Biol. 35: 1186–1197.10.1111/cobi.13663.

Christie, AP, Amano, T, Martin, PA, Petrovan, SO, Shackelford, GE, Simmons, BI, et al. 2021. The challenge of biased evidence in conservation. Conserv. Biol. 35: 249–262. 10.1111/cobi.13577.

Christie, AP, Amano, T, Martin, PA, Shackelford, GE, Simmons, BI, Sutherland, WJ 2019. Simple study designs in ecology produce inaccurate estimates of biodiversity responses. J. Appl. Ecol. 56: 2742–2754. 10.1111/1365-2664.13499.

Eisemberg, CC, Vogt, RC, Balestra, RAM, Reynolds, SJ, Christian, KA 2019. Don’t put all your eggs in one basket – Lessons learned from the largest-scale and longest-term wildlife conservation program in the Amazon Basin. Biol. Conserv. 238: 108182. 10.1016/j.biocon.2019.07.027.

Energisa 2024. Number of chelonian births in Tabuleiro do Guaporé drops from over 1.4 million in 2023 to 349,000 in 2024, (https://en.grupoenergisa.com.br/noticias/sustentabilidade/numero-de-nascimentos-de-quelonios-no-tabuleiro-do-guapore-reduz-de-mais). Accessed on 7 January 2025.

Forero-Medina, G, Ferrara, CR, Vogt, RC, Fagundes, CK, Balestra, RAM, Andrade, PCM, et al. 2021. On the future of the giant South American river turtle Podocnemis expansa. Oryx 55: 73–80. 10.1017/S0030605318001370.

Gibbs, JP, Amato, GD 2000. Turtle Conservation, edited by M. W. Klemens. Washington, DC: The Smithsonian Institution.

Greenland, S, Senn, SJ, Rothman, KJ, Carlin, JB, Poole, C, Goodman, SN, et al. 2016. Statistical tests, P values, confidence intervals, and power: a guide to misinterpretations. Eur. J. Epidemiol. 31: 337–350. 10.1007/s10654-016-0149-3.

Hallwass, G, Lopes, PF, Juras, AA, Silvano, RAM 2013. Fishers’ knowledge identifies environmental changes and fish abundance trends in impounded tropical rivers. Ecol. Appl. 23: 392–407. 10.1890/12-0429.1.

Halsey, LG 2019. The reign of the p-value is over: what alternative analyses could we employ to fill the power vacuum? Biol. Lett. 15: 20190174.10.1098/rsbl.2019.0174.

Hays, GC, Laloë, J-O, Seminoff, JA 2025. Status, trends and conservation of global sea turtle populations. Nature Reviews Biodiversity 1: 119–133. 10.1038/s44358-024-00011-y.

Hendrix, H, Pérez-Espona, S 2024. A Systematic Review of Population Monitoring Studies of Sea Turtles and Its Application to Conservation, Vol. 16, Diversity. 10.3390/d16030177.

Jones, JPG, Andriamarovololona, MM, Hockley, N, Gibbons, JM, Milner-Gulland, EJ 2008. Testing the use of interviews as a tool for monitoring trends in the harvesting of wild species. J. Appl. Ecol. 45: 1205–1212. 10.1111/j.1365-2664.2008.01487.x.

Keith, D, Akçakaya, HR, Butchart, SHM, Collen, B, Dulvy, NK, Holmes, EE, et al. 2015. Temporal correlations in population trends: Conservation implications from time-series analysis of diverse animal taxa. Biol. Conserv. 192: 247–257. 10.1016/j.biocon.2015.09.021.

Lacava, RV, Carvalho, DCdO, Pezzuti, JCB, da Silva, LCF, Miorando, PS, Fonseca, RA 2024. Recovery of the Giant South American River Turtle in four decades of a network-based conservation program in the Brazilian Amazon. Biodivers. Conserv. 10.1007/s10531-024-02971-1.

Lahoz-Monfort, JJ, Magrath, MJL 2021. A Comprehensive Overview of Technologies for Species and Habitat Monitoring and Conservation. Bioscience 71: 1038–1062. 10.1093/biosci/biab073.

Mogollones, SC, Rodríguez, DJ, Hernández, O, Barreto, GR 2010. A Demographic Study of the Arrau Turtle (Podocnemis expansa) in the Middle Orinoco River, Venezuela. Chelonian Conserv. Biol. 9: 79–89. 10.2744/CCB-0778.1.

Moll, D, Moll, EO 2004. The ecology, exploitation and conservation of river turtles, 1st ed. New York: Oxford University Press.

Muff, S, Nilsen, EB, O’Hara, RB, Nater, CR 2022. Rewriting results sections in the language of evidence. Trends Ecol. Evol. 37: 203–210. 10.1016/j.tree.2021.10.009.

Páez, VP, Lipman, A, Bock, BC, Heppell, SS 2015. A plea to redirect and evaluate conservation programs for South America’s Podocnemidid River Turtles. Chelonian Conserv. Biol. 14: 205–216. 10.2744/CCB-1122.1.

Peres, CA 2000. Effects of Subsistence Hunting on Vertebrate Community Structure in Amazonian Forests. Conserv. Biol. 14: 240–253. 10.1046/j.1523-1739.2000.98485.x.

Piacenza, SE, Richards, PM, Heppell, SS 2019. Fathoming sea turtles: monitoring strategy evaluation to improve conservation status assessments. Ecol. Appl. 29: e01942. 10.1002/eap.1942.

Wauchope, HS, Amano, T, Geldmann, J, Johnston, A, Simmons, BI, Sutherland, WJ, et al. 2021. Evaluating Impact Using Time-Series Data. Trends Ecol. Evol. 36: 196–205. 10.1016/j.tree.2020.11.001.

Zuur, AF, Ieno, EN, Elphick, CS 2010. A protocol for data exploration to avoid common statistical problems. Methods in Ecology and Evolution 1: 3–14. 10.1111/j.2041-210X.2009.00001.x.

