## Supplemental Material for "Misleading monitoring: more hatchlings do not represent turtle population recovery"

S1

Darren Norris

### Data evaluation: R code and results

#### Preparation: Packages and data

```
# Load necessary packages
library(plyr)
library(readxl)
library(tidyverse)
library(broom)
library(scales)
library(nlme)
library(MASS)
library(glmmTMB)
```

Load excel file from the Supplemental Material: <https://static-content.springer.com/esm/art%3A10.1007%2Fs10531-024-02971-1/MediaObjects/10531_2024_2971_MOESM1_ESM.xlsx>

```
hatchlings_pqa <- read_excel("10531_2024_2971_MOESM1_ESM.xlsx", sheet = "Sheet1",
                      skip = 7, .name_repair = "universal") |>
  filter(!is.na(Year))

# Long format.
hatchlings_pqa_long <- hatchlings_pqa |>
  dplyr::select(!Total) |>
  pivot_longer(cols = !Year, names_to = "beach",
               values_to = "hatchlings")
# To follow the sequence in the published results
bsig <- c("TRO", "BRA", "JAV", "ARA", "MOR", "ABU", "EMB", "MCR", "GUA",
          "CAM", "WBU")
bnames <- c("Trombetas", "Branco", "Javaes", "Araguaia", "Mortes",
           "Abufari", "Embaubal", "Monte Cristo", "Guaporé",
           "Camaleões", "Walter Bury")
bnamessm <- c("Trombetas", "Branco", "Javaés", "Araguaia", "Mortes",
           "Abufari", "Embaubal", "Monte.Cristo", "Guaporé",
           "Camalões.Island", "Walter.Bury")
beach_order <- data.frame(aid = 1:length(bsig), beach_sigla = bsig,
                          beach_names_t1 = bnames,
                          beach_names_s2 = bnamessm)
```

#### Reproduce models

Here I try to reproduce the original Generalized Linear Model (GLM) results. This is impossible as authors fail to detail methods sufficiently. The published text is unclear on how the GLMs are modelled. They also fail to detail how they checked for the best model. Here I assume the authors “best model” was selected via AIC, and the results presented by authors in Figure 2 are from the negative binomial GLM model.

I use function glm.nb() to get estimates, AIC, theta and deviance explained.

```
# Coefficient estimates
models_tidy_glm_nb <- hatchlings_pqa_long |>
  filter(!is.na(hatchlings)) |>
  group_by(beach) |>
  group_modify(.f = ~ tidy(MASS::glm.nb(hatchlings ~ Year, data=.x)))

# Model summary
glance_glmnb <- function(x){

df <- na.omit(x)
# model
model_glmnb <- MASS::glm.nb(hatchlings ~ Year, data=df)

# Summary of the glm.nb model results
s_glmnb <- summary(model_glmnb)
mytheta <- s_glmnb$theta
myaic <- s_glmnb$aic
# Calculate the null deviance (deviance of the null model)
null_deviance <- s_glmnb$null.deviance
# Calculate the residual deviance (deviance of the fitted model)
residual_deviance <- s_glmnb$deviance
# Calculate the percentage deviance explained
deviance_explained <- 100 * ((null_deviance - residual_deviance) / null_deviance)

dfout <- data.frame(theta = mytheta, AIC = myaic,
                    dev_exp = round(deviance_explained, 1))
dfout
}

models_glance_glm_nb <- plyr::ddply(hatchlings_pqa_long, .(beach),
                                    .fun = glance_glmnb)
models_glance_glm_nb |>
  left_join(
models_tidy_glm_nb |>
  filter(term == "Year")
) |>
  arrange(estimate) |>
  knitr::kable("simple", digits = c(1, 1, 1, 1, 1, 3, 3, 2, 5))
Joining with `by = join_by(beach)`
```

Table 2: GLM-nb summary.

| beach | theta | AIC | dev_exp | term | estimate | std.error | statistic | p.value |
| --- | --- | --- | --- | --- | --- | --- | --- | --- |
| Trombetas | 1.4 | 1005.2 | 51.5 | Year | -0.073 | 0.011 | -6.62 | 0.00000 |
| Branco | 1.1 | 899.9 | 25.2 | Year | -0.057 | 0.012 | -4.90 | 0.00000 |
| Javaés | 2.8 | 597.6 | 14.3 | Year | -0.032 | 0.013 | -2.39 | 0.01682 |
| Mortes | 1.1 | 919.6 | 0.1 | Year | 0.003 | 0.015 | 0.23 | 0.81814 |
| Abufari | 2.9 | 1104.8 | 10.1 | Year | 0.019 | 0.007 | 2.59 | 0.00949 |
| Embaubal | 2.0 | 1119.9 | 12.3 | Year | 0.022 | 0.009 | 2.45 | 0.01443 |
| Araguaia | 0.6 | 915.8 | 3.7 | Year | 0.025 | 0.021 | 1.18 | 0.23615 |
| Monte.Cristo | 2.0 | 1182.9 | 56.6 | Year | 0.083 | 0.008 | 9.93 | 0.00000 |
| Camalões.Island | 1.4 | 935.9 | 58.4 | Year | 0.095 | 0.011 | 8.74 | 0.00000 |
| Walter.Bury | 3.7 | 941.9 | 85.0 | Year | 0.118 | 0.007 | 16.48 | 0.00000 |
| Guaporé | 3.4 | 846.6 | 84.5 | Year | 0.127 | 0.008 | 15.74 | 0.00000 |

In the article authors identify six areas where “populations increased over time with statistical significance”. The six areas were Abufari and Embaubal (“more subtle increase”), and Monte Cristo, Guaporé, Camaleões, and Walter Bury with “very evident increase of hatchlings released over the 40 years.”. The results presented in Table 2 closely match those reported by authors.

A previous study reported increases in 4 of 11 PQA areas (Eisemberg et al. (2019)). Another study reported increases in 4 of 9 PQA areas (Forero-Medina et al. (2019)). In the Discussion authors attribute the more positive results compared to previous studies to the increased number of monitoring years: ” Based on a more extended data set than previous studies on the Brazilian PQA public dataset (Eisemberg et al. 2019; Forero-Medina et al. 2019), our results show a far more positive trend of increase over 43 years in most areas and for all the nesting sites pooled together. ”

I can test this by running models with a subset of the authors data that includes only the 30 years used in the earlier studies.

```
hatchlings_pqa_sub <- hatchlings_pqa_long |>
  filter(Year <= 2008)
# Coefficient estimates
models_tidy_glm_nb <- hatchlings_pqa_sub |>
  filter(!is.na(hatchlings)) |>
  group_by(beach) |>
  group_modify(.f = ~ tidy(MASS::glm.nb(hatchlings ~ Year, data=.x)))

# Model summary
glance_glmnb <- function(x){

df <- na.omit(x)
# model
model_glmnb <- MASS::glm.nb(hatchlings ~ Year, data=df)

# Summary of the glm.nb model results
s_glmnb <- summary(model_glmnb)
mytheta <- s_glmnb$theta
myaic <- s_glmnb$aic
# Calculate the null deviance (deviance of the null model)
null_deviance <- s_glmnb$null.deviance
# Calculate the residual deviance (deviance of the fitted model)
residual_deviance <- s_glmnb$deviance
# Calculate the percentage deviance explained
deviance_explained <- 100 * ((null_deviance - residual_deviance) / null_deviance)

dfout <- data.frame(theta = mytheta, AIC = myaic,
                    dev_exp = round(deviance_explained, 1))
dfout
}

models_glance_glm_nb <- plyr::ddply(hatchlings_pqa_sub, .(beach),
                                    .fun = glance_glmnb)
models_glance_glm_nb |>
  left_join(
models_tidy_glm_nb |>
  filter(term == "Year")
) |>
  arrange(estimate) |>
  knitr::kable("simple", digits = c(1, 1, 1, 1, 1, 3, 3, 2, 5))
Joining with `by = join_by(beach)`
```

Table 3: GLM-nb summary. Results from 1997 - 2008 subset.

| beach | theta | AIC | Dev exp | term | estimate | std. Error | statistic | P-value |
| --- | --- | --- | --- | --- | --- | --- | --- | --- |
| Trombetas | 1.5 | 708.1 | 48.0 | Year | -0.108 | 0.019 | -5.60 | 0.00000 |
| Araguaia | 0.9 | 571.2 | 5.1 | Year | -0.057 | 0.034 | -1.67 | 0.09432 |
| Javaés | 4.4 | 503.8 | 5.1 | Year | -0.017 | 0.014 | -1.20 | 0.23021 |
| Mortes | 1.3 | 631.2 | 0.0 | Year | -0.003 | 0.025 | -0.13 | 0.89799 |
| Embaubal | 1.4 | 733.1 | 2.5 | Year | 0.016 | 0.019 | 0.82 | 0.41029 |
| Branco | 5.5 | 635.0 | 8.3 | Year | 0.021 | 0.012 | 1.78 | 0.07574 |
| Abufari | 2.8 | 743.3 | 29.3 | Year | 0.052 | 0.013 | 3.90 | 0.00009 |
| Camalões.Island | 1.1 | 587.3 | 23.5 | Year | 0.080 | 0.022 | 3.65 | 0.00026 |
| Guaporé | 4.2 | 550.4 | 77.5 | Year | 0.121 | 0.012 | 9.69 | 0.00000 |
| Walter.Bury | 3.7 | 597.5 | 76.1 | Year | 0.121 | 0.013 | 9.41 | 0.00000 |
| Monte.Cristo | 6.0 | 750.5 | 84.2 | Year | 0.127 | 0.009 | 14.72 | 0.00000 |

As expected, there are differences over time. But the differences do not follow authors’ affirmation that the more positive outcomes come from the longer data set. For example, the counts at Branco have worsened over time. Branco had a negative trend with the full data set as reported in Table 1 by Lacava et. al. (Table 2), but running the same analysis with the 30 year subset Branco had a statistically insignificant positive increase in hatchlings released (Table 3). The more positive outcome overall among beaches is a result of a single beach - Embaubal. Embaubal changes from a significantly insignificant increase in the shorter time series, to a statistically significant increase in the full time series (Table 3, Table 2). Simply put the situation can worsen or improve over time.

The differences in the results presented by Lacava et al. (2024) most likely come from the far more robust analysis presented in the earlier studies. The earlier studies used a combination of indicators not only hatchling counts and also attempted to correct for methodological differences over time (Eisemberg et al. (2019), Forero-Medina et al. (2019)). As such the more cautious conclusions presented in the earlier studies appear to have much stronger support.

#### Significance testing

I first compare the distribution of the response variable across the different areas.

```
# check distribution
hatchlings_pqa_long |>
  filter(!is.na(hatchlings)) |>
ggplot(aes(x = hatchlings)) +
  geom_histogram(fill = "blue", color = "black", bins = 6) +
  labs(title = "Histogram of hatchlings released",
       x = "Hatchlings released (thousands)", y = "Frequency") +
  scale_x_continuous(labels = label_number(scale = 1/1000)) +
  facet_wrap(~beach, scales = "free")
```

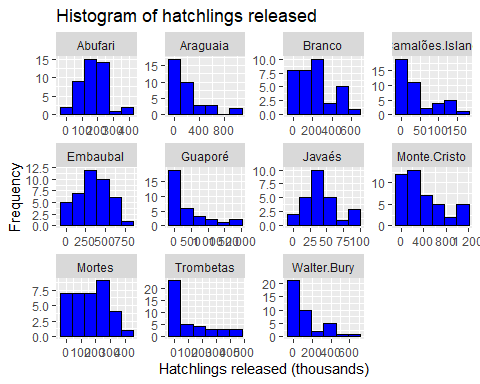

```
# Perform Shapiro-Wilk test for each group
# Approximately normal
hatchlings_pqa_long |>
  filter(!is.na(hatchlings)) |>
  group_by(beach) |>
  summarize(shapiro_test = list(shapiro.test(hatchlings))) |>
  mutate(tidy_shapiro = map(shapiro_test, broom::tidy)) |>
  unnest(tidy_shapiro)
# A tibble: 11 × 5
   beach           shapiro_test statistic     p.value method
   <chr>           <list>           <dbl>       <dbl> <chr>
 1 Abufari         <htest>          0.961 0.151       Shapiro-Wilk normality te…
 2 Araguaia        <htest>          0.766 0.00000464  Shapiro-Wilk normality te…
 3 Branco          <htest>          0.938 0.0523      Shapiro-Wilk normality te…
 4 Camalões.Island <htest>          0.772 0.00000118  Shapiro-Wilk normality te…
 5 Embaubal        <htest>          0.973 0.434       Shapiro-Wilk normality te…
 6 Guaporé         <htest>          0.729 0.00000182  Shapiro-Wilk normality te…
 7 Javaés          <htest>          0.919 0.0436      Shapiro-Wilk normality te…
 8 Monte.Cristo    <htest>          0.885 0.000397    Shapiro-Wilk normality te…
 9 Mortes          <htest>          0.953 0.136       Shapiro-Wilk normality te…
10 Trombetas       <htest>          0.712 0.000000117 Shapiro-Wilk normality te…
11 Walter.Bury     <htest>          0.756 0.000000907 Shapiro-Wilk normality te…
# skewness for each group
hatchlings_pqa_long |>
  filter(!is.na(hatchlings)) |>
  group_by(beach) |>
  summarize(skew = list(moments::skewness(hatchlings))) |>
  mutate(tidy_skew = map(skew, broom::tidy)) |>
  unnest(tidy_skew)
# A tibble: 11 × 3
   beach           skew           x
   <chr>           <list>     <dbl>
 1 Abufari         <dbl [1]> 0.296
 2 Araguaia        <dbl [1]> 1.76
 3 Branco          <dbl [1]> 0.517
 4 Camalões.Island <dbl [1]> 1.16
 5 Embaubal        <dbl [1]> 0.0101
 6 Guaporé         <dbl [1]> 1.65
 7 Javaés          <dbl [1]> 0.764
 8 Monte.Cristo    <dbl [1]> 0.806
 9 Mortes          <dbl [1]> 0.100
10 Trombetas       <dbl [1]> 1.32
11 Walter.Bury     <dbl [1]> 1.41
```

Preliminary descriptive analysis shows differences in the distribution of responses among beaches.

##### Temporal autocorrelation

As the analysis is based on counts over time, temporal autocorrelation needs to be explicitly included in the models. There are several reasons why this is important but generally the most important reason is to avoid type I errors (Carroll and Pearson (2000), Keith et al. (2015), Wolkovich et al. (2014), Zuur, Ieno, and Elphick (2009)).

Here I include temporal autocorrelation using Generalized Linear Mixed Models with the package glmmTMB (Brooks et al. (2017)).

First identify which distribution family is most appropriate for each area.

```
# Model summaries to identify best distribution family
glance_glmmTMB <- function(x){
df_ab <- na.omit(x)
# Convert year to a factor
df_ab$Yearf <- factor(df_ab$Year)
# Create a single group factor
df_ab$group <- factor(rep(1, nrow(df_ab)))

model_glmmTMB_gau <- glmmTMB::glmmTMB(hatchlings ~ Year + ar1(0 + Yearf | group),
                data = df_ab)
model_glmmTMB_nb <- glmmTMB::glmmTMB(hatchlings ~ Year + ar1(0 + Yearf | group),
                data = df_ab,
                family = nbinom1)
model_glmmTMB_po <- glmmTMB::glmmTMB(hatchlings ~ Year + ar1(0 + Yearf | group),
                data = df_ab,
                family = poisson)

s_glmmTMB_gau <- summary(model_glmmTMB_gau)
s_glmmTMB_gau <- data.frame(values = s_glmmTMB_gau$AICtab)
s_glmmTMB_gau$terms <- row.names(s_glmmTMB_gau)
s_glmmTMB_gau$aid <- factor("a")
s_glmmTMB_gau <- s_glmmTMB_gau |> pivot_wider(id_cols = aid,
                                names_from = terms, values_from = values)


s_glmmTMB_nb <- summary(model_glmmTMB_nb)
s_glmmTMB_nb <- data.frame(values = s_glmmTMB_nb$AICtab)
s_glmmTMB_nb$terms <- row.names(s_glmmTMB_nb)
s_glmmTMB_nb$aid <- factor("a")
s_glmmTMB_nb <- s_glmmTMB_nb |> pivot_wider(id_cols = aid,
                                names_from = terms, values_from = values)
s_glmmTMB_po <- summary(model_glmmTMB_po)
s_glmmTMB_po <- data.frame(values = s_glmmTMB_po$AICtab)
s_glmmTMB_po$terms <- row.names(s_glmmTMB_po)
s_glmmTMB_po$aid <- factor("a")
s_glmmTMB_po <- s_glmmTMB_po |> pivot_wider(id_cols = aid,
                                names_from = terms, values_from = values)
dfout <- bind_rows(
  s_glmmTMB_gau |> dplyr::select(!aid) |>
  mutate(model_family = "gaussian"),
  s_glmmTMB_nb |> dplyr::select(!aid) |>
  mutate(model_family = "negative binomial"),
 s_glmmTMB_po |> dplyr::select(!aid) |>
  mutate(model_family = "poisson")) |>
  relocate(model_family)

dfout
}

models_glance_glmmTMB <- plyr::ddply(hatchlings_pqa_long, .(beach),
                                    .fun = glance_glmmTMB)
models_glance_glmmTMB <- models_glance_glmmTMB |>
  group_by(beach) |> arrange(AIC) |>
  mutate(delta_aic = AIC - min(AIC, na.rm = TRUE)) |>
  ungroup() |> arrange(beach, AIC)

models_glance_glmmTMB |>
  filter(delta_aic == 0) |>
  knitr::kable()
```

Table 4: Generalized Linear Mixed Model summary.

| **beach** | **Model family** | **AIC** | **BIC** | **logLik** | **deviance** | **df.resid** | **delta_aic** |
| --- | --- | --- | --- | --- | --- | --- | --- |
| Abufari | negative binomial | 1098.1 | 1106.9 | -544.1 | 1088.1 | 38 | 0 |
| Araguaia | negative binomial | 917.9 | 925.6 | -453.9 | 907.9 | 30 | 0 |
| Branco | negative binomial | 866.6 | 874.2 | -428.3 | 856.6 | 29 | 0 |
| Camalões.Island | negative binomial | 925.5 | 934.2 | -457.7 | 915.5 | 37 | 0 |
| Embaubal | negative binomial | 1116.8 | 1125.3 | -553.4 | 1106.8 | 36 | 0 |
| Guaporé | poisson | 841.8 | 847.8 | -416.9 | 833.8 | 29 | 0 |
| Javaés | poisson | 602.2 | 607.2 | -297.1 | 594.2 | 22 | 0 |
| Monte.Cristo | negative binomial | 1199.0 | 1207.9 | -594.5 | 1189.0 | 39 | 0 |
| Mortes | poisson | 935.9 | 942.1 | -463.9 | 927.9 | 31 | 0 |
| Trombetas | negative binomial | 1013.8 | 1022.4 | -501.9 | 1003.8 | 36 | 0 |
| Walter.Bury | negative binomial | 944.3 | 952.8 | -467.2 | 934.3 | 35 | 0 |

Now run the Generalized Linear Mixed Models including temporal autocorrelation.

```
# Predictions using best of negative binomial/poisson.
dfin <- hatchlings_pqa_long |>
  left_join(models_glance_glmmTMB |>
              filter(delta_aic == 0) |>
              dplyr::select(beach, model_family))

# Function to run glmmTMB and get predictions.
pred_beaches_glmmTMB <- function(x) {
df_ab <- na.omit(x)
# Convert year to a factor
df_ab$Yearf <- factor(df_ab$Year)
# Create a single group factor
df_ab$group <- factor(rep("a", nrow(df_ab)))
myfamily <- x$model_family[1]
if(myfamily == "negative binomial"){
model_glmmTMB <- glmmTMB::glmmTMB(hatchlings ~ Year + ar1(0 + Yearf | group),
                data = df_ab,
                family = nbinom1)
}else{
  model_glmmTMB <- glmmTMB::glmmTMB(hatchlings ~ Year + ar1(0 + Yearf | group),
                data = df_ab,
                family = poisson)
}
dfnew_ab <- data.frame(Year = seq(min(df_ab$Year),
                                  max(df_ab$Year), by = 1))
# Convert year to a factor
dfnew_ab$Yearf <- factor(dfnew_ab$Year)
# Create a single group factor
dfnew_ab$group <- factor(rep("a", nrow(dfnew_ab)))
p_tmb <- predict(model_glmmTMB, newdata = dfnew_ab, type = "response",
              se.fit = TRUE, allow.new.levels=TRUE)
dfnew_ab <- dfnew_ab |>
  dplyr::mutate(glmmTMB_fit = p_tmb$fit,
         glmmTMB_se = p_tmb$se.fit) |>
  dplyr::mutate(lcl  = glmmTMB_fit - (1.96 * glmmTMB_se),
         ucl = glmmTMB_fit + (1.96 * glmmTMB_se))

dfres <- dfnew_ab |> data.frame()
dfres
}

# Run function for each beach.
model_res_glmmTMB <- plyr::ddply(dfin, c("beach"),
                         .fun = pred_beaches_glmmTMB)
```

Now plot predictions.

```
beach_order$aidf <- factor(beach_order$aid)
levels(beach_order$aidf) <- beach_order$beach_names_t1

# Figure
fig_hatch_glmmTMB <- hatchlings_pqa_long |>
  left_join(model_res_glmmTMB) |>
  left_join(beach_order, join_by(beach == beach_names_s2)) |>
  mutate(lcl = ifelse(lcl < 0 , 0, lcl)) |>
  ggplot(aes(x = Year, y = hatchlings)) +
  geom_point() +
  geom_ribbon(aes(ymin = lcl, ymax = ucl), alpha=0.2) +
  geom_line(aes(x = Year, y = glmmTMB_fit), colour = "blue") +
  scale_y_continuous(labels = label_number(scale = 1/1000)) +
  facet_wrap( ~ aidf, scales = "free_y", ncol = 3) +
  theme_bw() +
  labs(y = "Hatchlings released (thousands)")

fig_hatch_glmmTMB
```

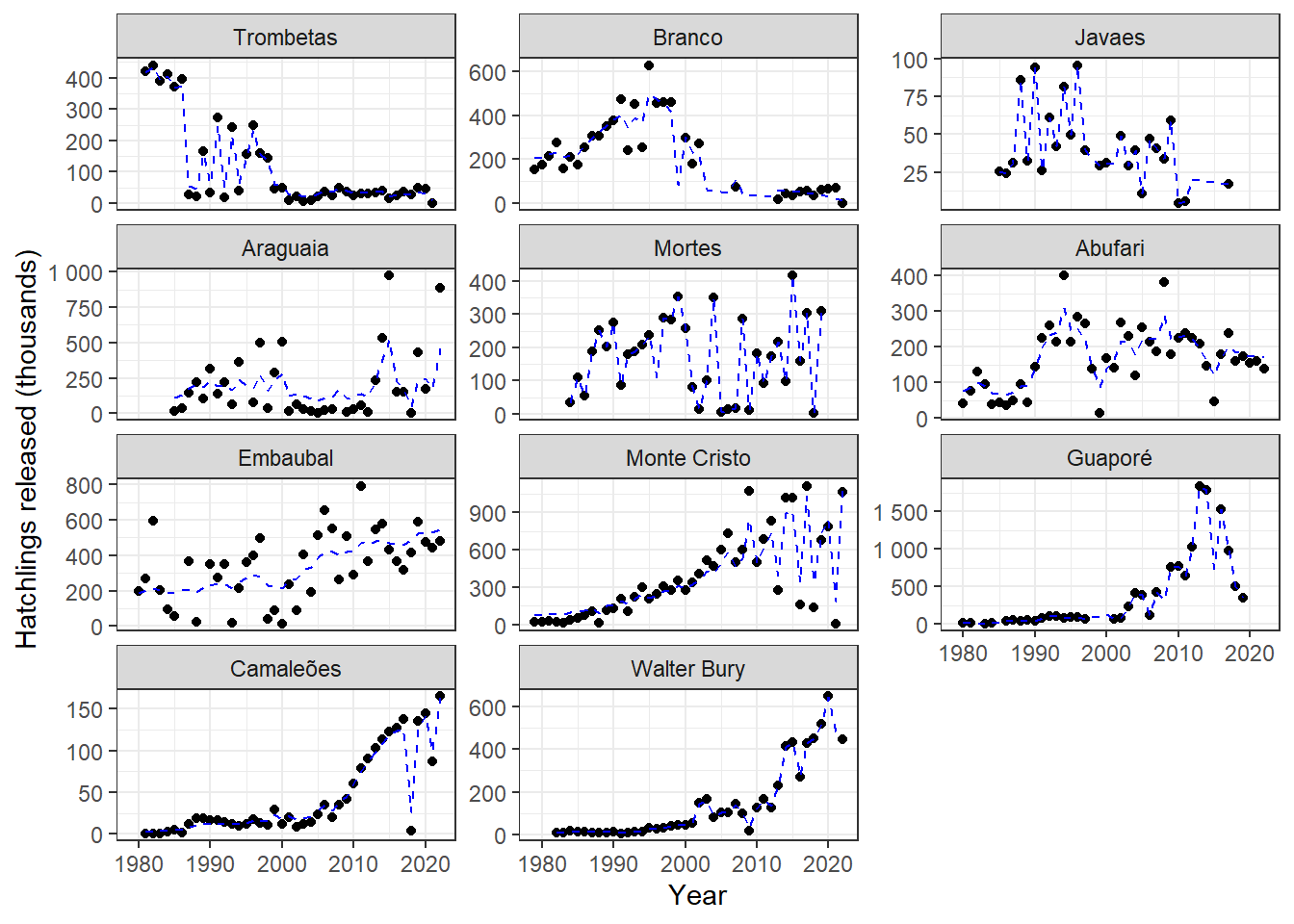

Figure 2: The number of Giant South American River Turtle (Podocnemis expansa) hatchlings released from 11 reproductive areas monitored by the Brazilian turtle conservation program (PQA). Dashed blue lines show predictions from Generalized Linear Mixed Model including temporal autocorrelation.

Now obtain the Generalized Linear Mixed Model coefficients.

```
tidy_glmmTMB2 <- function(x){
df_ab <- na.omit(x)
# Convert year to a factor (this is still needed)
df_ab$Yearf <- factor(df_ab$Year)
# Create a single group factor
df_ab$group <- factor(rep(1, nrow(df_ab)))
myfamily <- x$model_family[1]
if(myfamily == "negative binomial"){
model_glmmTMB <- glmmTMB::glmmTMB(hatchlings ~ Year + ar1(0 + Yearf | group),
                data = df_ab,
                family = nbinom1)
}else{
  model_glmmTMB <- glmmTMB::glmmTMB(hatchlings ~ Year + ar1(0 + Yearf | group),
                data = df_ab,
                family = poisson)
}

s_glmmTMB <- summary(model_glmmTMB)
dfout <- data.frame(s_glmmTMB$coefficients$cond)
dfout$term <- row.names(dfout)
row.names(dfout) <- NULL
dfout
}

models_tidy_glmmTMB_final <- plyr::ddply(dfin, .(beach),
                                    .fun = tidy_glmmTMB2)
models_tidy_glmmTMB_final |>
  filter(term == "Year") |>
  arrange(Estimate) |>
  knitr::kable("simple", digits = c(0, 3, 3, 2, 5, 0))
```

Table 5: Model coefficients from Generalized Linear Mixed Models accounting for temporal autocorrelation.

| beach | Estimate | Std Error | Z-value | P-value | term |
| --- | --- | --- | --- | --- | --- |
| Branco | -0.079 | 0.027 | -2.96 | 0.00311 | Year |
| Trombetas | -0.075 | 0.013 | -5.56 | 0.00000 | Year |
| Javaés | -0.037 | 0.016 | -2.33 | 0.02004 | Year |
| Mortes | -0.020 | 0.021 | -0.97 | 0.33312 | Year |
| Araguaia | 0.006 | 0.021 | 0.27 | 0.78416 | Year |
| Abufari | 0.020 | 0.012 | 1.68 | 0.09202 | Year |
| Embaubal | 0.028 | 0.009 | 3.00 | 0.00269 | Year |
| Monte.Cristo | 0.055 | 0.011 | 4.82 | 0.00000 | Year |
| Camalões.Island | 0.092 | 0.011 | 8.67 | 0.00000 | Year |
| Guaporé | 0.112 | 0.017 | 6.45 | 0.00000 | Year |
| Walter.Bury | 0.118 | 0.011 | 11.03 | 0.00000 | Year |

Once temporal autocorrelation is controlled for there are “statistically significant” increases in the number of hatchlings released at five of the 11 nesting areas (Table 5). Abufari beach is no longer “statistically significant”. This area had a jump in the number of hatchlings released around 1990 (Figure 2). In the early years there was an average of approximately 50 thousand hatchlings released until 1990, then after 1990 a jump to approximately 200,000 hatchlings released. Such a sudden and large change is not consistent with population dynamics of turtle species and was not be captured by the simplistic GLM approach adopted by the authors. The most likely explanation for such a jump is a change in management such as increasing the beach area protected, change in counting technique etc.

### References

Baker, Monya. 2016. “Statisticians Issue Warning over Misuse of P Values.” *Nature* 531 (7593): 151–51. <https://doi.org/10.1038/nature.2016.19503>.

Brooks, Mollie,E., Kasper Kristensen, Koen,J.,van Benthem, Arni Magnusson, Casper,W. Berg, Anders Nielsen, Hans,J. Skaug, Martin Mächler, and Benjamin,M. Bolker. 2017. “glmmTMB Balances Speed and Flexibility Among Packages for Zero-Inflated Generalized Linear Mixed Modeling.” *The R Journal* 9 (2): 378. <https://doi.org/10.32614/rj-2017-066>.

Carroll, Steven S., and David L. Pearson. 2000. “Detecting and Modeling Spatial and Temporal Dependence in Conservation Biology.” *Conservation Biology* 14 (6): 1893–97. <https://doi.org/10.1111/j.1523-1739.2000.99432.x>.

Christie, Alec P., Tatsuya Amano, Philip A. Martin, Gorm E. Shackelford, Benno I. Simmons, and William J. Sutherland. 2019. “Simple Study Designs in Ecology Produce Inaccurate Estimates of Biodiversity Responses.” Edited by Júlio Louzada. *Journal of Applied Ecology* 56 (12): 2742–54. <https://doi.org/10.1111/1365-2664.13499>.

Eisemberg, C. C., R. C. Vogt, R. A. M. Balestra, S. J. Reynolds, and K. A. Christian. 2019. “Don’t Put All Your Eggs in One Basket – Lessons Learned from the Largest-Scale and Longest-Term Wildlife Conservation Program in the Amazon Basin.” *Biological Conservation* 238: 108182. <https://doi.org/10.1016/j.biocon.2019.07.027>.

Forero-Medina, German, Camila R. Ferrara, Richard C. Vogt, Camila K. Fagundes, Rafael Antônio M. Balestra, Paulo C. M. Andrade, Roberto Lacava, et al. 2019. “On the Future of the Giant South American River Turtle *Podocnemis Expansa*.” *Oryx* 55 (1): 73–80. <https://doi.org/10.1017/s0030605318001370>.

Gopalaswamy, Arjun M., Nicholas B. Elliot, Shadrack Ngene, Femke Broekhuis, Alexander Braczkowski, Peter Lindsey, Craig Packer, and Nils Chr. Stenseth. 2022. “How “Science” Can Facilitate the Politicization of Charismatic Megafauna Counts.” *Proceedings of the National Academy of Sciences* 119 (20). <https://doi.org/10.1073/pnas.2203244119>.

Halsey, Lewis G. 2019. “The Reign of the *p* -Value Is over: What Alternative Analyses Could We Employ to Fill the Power Vacuum?” *Biology Letters* 15 (5): 20190174. <https://doi.org/10.1098/rsbl.2019.0174>.

Keith, David, H. Resit Akçakaya, Stuart H. M. Butchart, Ben Collen, Nicholas K. Dulvy, Elizabeth E. Holmes, Jeffrey A. Hutchings, et al. 2015. “Temporal Correlations in Population Trends: Conservation Implications from Time-Series Analysis of Diverse Animal Taxa.” *Biological Conservation* 192 (December): 247–57. <https://doi.org/10.1016/j.biocon.2015.09.021>.

Lacava, Roberto Victor, Dennison Célio de Oliveira Carvalho, Juarez Carlos Brito Pezzuti, Leo Caetano Fernandes da Silva, Priscila Saikoski Miorando, and Raphael Alves Fonseca. 2024. “Recovery of the Giant South American River Turtle in Four Decades of a Network-Based Conservation Program in the Brazilian Amazon.” *Biodiversity and Conservation*, November. <https://doi.org/10.1007/s10531-024-02971-1>.

Muff, Stefanie, Erlend B. Nilsen, Robert B. O’Hara, and Chloé R. Nater. 2022. “Rewriting Results Sections in the Language of Evidence.” *Trends in Ecology & Evolution* 37 (3): 203–10. <https://doi.org/10.1016/j.tree.2021.10.009>.

Pauly, Daniel, Villy Christensen, Sylvie Guénette, Tony J. Pitcher, U. Rashid Sumaila, Carl J. Walters, R. Watson, and Dirk Zeller. 2002. “Towards Sustainability in World Fisheries.” *Nature* 418 (6898): 689–95. <https://doi.org/10.1038/nature01017>.

Wauchope, Hannah S., Tatsuya Amano, Jonas Geldmann, Alison Johnston, Benno I. Simmons, William J. Sutherland, and Julia P. G. Jones. 2021. “Evaluating Impact Using Time-Series Data.” *Trends in Ecology & Evolution* 36 (3): 196–205. <https://doi.org/10.1016/j.tree.2020.11.001>.

Wolkovich, E. M., B. I. Cook, K. K. McLauchlan, and T. J. Davies. 2014. “Temporal Ecology in the Anthropocene.” Edited by Franck Courchamp. *Ecology Letters* 17 (11): 1365–79. <https://doi.org/10.1111/ele.12353>.

Zuur, Alain F., Elena N. Ieno, and Chris S. Elphick. 2009. “A Protocol for Data Exploration to Avoid Common Statistical Problems.” *Methods in Ecology and Evolution* 1 (1): 3–14. <https://doi.org/10.1111/j.2041-210x.2009.00001.x>.
